# Prevalence of Bovine *Cysticercosis*, Human Taeniasis and Asociated Risk Factors in Selected Towns of Silte Zone, Southern Ethiopia

**DOI:** 10.1101/2023.01.25.525623

**Authors:** Solomon musema mussa

## Abstract

Bovine cysticercosis is an infection of cattle caused by the larval stage of Cysticercus Bovis, the human intestinal cestodes and it has economic important and public health importance. Across-sectional study was conducted from March 2022 to August 2022 to determine the prevalence of Bovine Cysticercosis, Human Taeniasis and associated risk factors in selected towns of Silte zone. The study was conducted in Silte zone municipal abbatoir, Southern Ethiopia in cattle slaughtered randomly selected towns,namely, at Kilto,Kutery,Kirate and Werabe Tawon municipal abattoirs. Systematic random sampling technique was used to collect all the necessary data from abattoir survey of the study animals and questionnaire survey collected data stored by using in Epi data software and analyzed using (SPSS-22) software program. The sample size required for this study was determined to be based on the expected prevalence (50%) of C. bovis and the 5% desired absolute precision and 95% CI. Out of 422 carcasses examined during the study period in 4 municipality abattoirs, 36 % were infected with C.bovis. A prevalence of 35.7%, 35.92%, 37.96% and 34.11% were Kilto,Kutery,Kirate and Werabe Tawon were observed, respectively. Among considered risk factors, sex, body condition and breed showed statistically significant difference in prevalence of C. bovis in study area. Regarding the anatomical distribution of the cysts, shoulder muscle showed highest prevalence 28.44%, was followed by tongue 20.79% tricepsbrachi muscle 18.35%, masseter muscle 17.74%, heart 12.84%, and Liver 1.8%,. Out of 110 respondents, 64.5% had contracted T.saginata. Among considered risk factors, sex, age, occupation, education status, raw meat consumption and knowledge about the disease showed statistically significant difference in prevalence of T.saginata in study area. The findings of this study indicated that importance of Bovine Cysticercosis and Taeniasis in the study area. Therefore, attention should be given to the public awareness, detailed meat inspection to be safe to public health, consumption of well-cooked cattle meat should be implemented for breaking the cycle of the diseases and promote meat industry in the country.

## 1. INTRODUCTION

Ethiopia has one of the largest inventories in Africa with livestock currently supporting and sustaining the livelihoods of an estimated 80 % of the rural poor (refereciaed). Animal rearing is an integral part of the agricultural production and estimated livestock population is 70 million cattle, 42.9 million sheep and 52.5 million goats (CSA, 2021/22). Livestock are the main stays of the livelihood of the majority of the human population by giving draft power, income to farming communities, means of investment and important source of foreign exchange earning to the nation. Moreover, livestock are important cultural resources, social safety nets and means of saving, and also supply for crop production and transports (DACA, 2006). In Ethiopia, the livestock sector contributes about 30% of the agricultural GDP and 19% to the export earnings (CSA, 2021/22). In Sub-Saharan Africa, livestock diseases, negatively affect the public health and impede economic growth by incurring direct (morbidity, mortality) and indirect economic losses (Bekele et al., 2010). Among highly prevalent diseases, bovine cysticercosis is most common and it causes that economic losses, and public health problems (Tariku and Beredo, 2019). It is a muscular infection of cattle caused by the larvae of the human intestinal cestodes (Soulsby, 2009).

Bovine cysticercosis is a food borne disease caused by Taenia saginata with humans as the final host and cattle as the intermediate host. Infection of human by Taenia saginata occurs through ingestion of raw or undercooked meat containing Cysticercus Bovis while; infection of cattle with Cysticercus Bovis occurs through ingestion of Taenia saginata eggs. The parasite population of these species consists of three distinct sub populations: adult Tapeworms in the definitive host (man), larvae (Cysticercus or metacestodes) in the intermediate host (pigs or cattle), and eggs in the environment (Dupuy et al., 2014.).

T. saginata is global distributed in both developed and developing countries. However, it is high reported incidence cases in Africa when compared with other parts of the world (Abunna et al., 2007).This parasite epidemiology is ethnically and culturally determined with estimation annually cases and mortality rate around 50-77 million and 50, 000 people respectively (Birhanu et al., 2018). T. saginata which is known as beef tapeworm has two stages of development, intermediate host and final host in cattle and human respectively. Larval stage of this parasite occurs in heart and skeletal muscles of intermediate host and adult worm locates in intestine of final host. Visual inspection of carcass and other organs is the primary detection method of Bovine Cysticercosis because commonly found area of carcass are the external and internal masseter pterygoid muscles, heart, tongue, diaphragm, and esophagus (Oladele et al., 2004).

In discriminate defecation, due to lacking latrine facilities, is common practice especially by the rural community in Ethiopia where more than 80% of the populations reside. The common traditional animal husbandry practices in Ethiopia (free grazing in cattle) mainly allow free access of cattle to the contaminated environment and perpetuate transmission of Cysticercosis, due to the fact that cattle become infected by ingestion of pasture/feed or water contaminated with T. saginata eggs (Kifle et al., 2015). It is associated with poor hygiene and local factors including cultural background, such as eating meat without proper cooking (raw), economic condition and religious beliefs, close proximity of humans to cattle kept with little or no distinction between companion or utility functions (Fralova, 2014).

As reported in Ethiopia from several authors, prevalence the average 30% from different abattoirs in the country (Adugna et al., 2013) and 30.7% at Eastern Shoa of Oromia (Abdela et al., 2019). Taenia saginata taeniasis/cysticercosis is high economic and public health impacts in Ethiopia; as a result control and prevention of the disease has great importance. One of the prerequisite for implementing control and prevention action is information on prevalence and associated risk factors in Silte Zone esepechaialy in Kilto, Kutery, Kirate and Werabe town there is limited work that indicates the status, risk factors and public health importance of T. saginata taeniasis/ Cysticercosis. Therfor the objective of this study is to determine the prevalence of Bovine Cysticercosis, Human Taeniasis and identify potential risk factors of the disease in study area.

## 2. Materials and Methods

### 2.1. Description of Study Area

Silte zone was one of the 15 zones in SNNPRS. The zone was sub divided into 10 Woreda and 03 town administration. The study was conducted in Werabe capital town of Silte Zone and other 4 districts, namely, Misrak Azernet, Hulbareg, Alicho Weriro, and Werabe which are head towen,of Kilto,Kutery,Kirate and Werabe districts respectively. Silte is bordered on the south by Halaba Zone, on the south west by Hadiya, on the north by Gurage, and on the east by the Oromia Region. Werabe was the capital tawon of Silte zone, which is located 175km South of Addis Ababa and away from 222 km from Hawassa.

Agro-ecologically, Silte zone is classified as low land, mid land and high land and its climate fell under the category of mid land by 73.5 percent, high land by 23.01 percent and low land by 3.4 percent. It is located in between 7.43° - 8.10°N latitude and 37.86° – 38.53°E longitudes. The elevation varies from 1,501 meters to 2,500 meters above sea level. The average annual rainfall is 800-1818 milliliters while the average annual temperature ranges from 20.1°C to 25°C. The land utilization is 65.84% cultivated land, 8.78% grazing land, 4.27% forest bushes, shrub land 3.97% and 17.14% were covered by others (Silte Zone Agricultural Office, 2020).

### 2.2. Study Population

The study animals were cattle brought from different parts of the surrounding areas for slaughtering purpose in selected abattoirs Tawons of Silte zone. The study population consists of cattle at different ages, sex, origins and breeds categories in the study area. For questionnaire survey the target population was people living in Misrak Azernet, Hulbareg, Alicho Weriro, and Werabe towns.

### 2.3. Study Design

A cross-sectional study design was conducted in abattoir selected Tawons of Silte zone from March 2022 to August 2022 to determine the prevalence of Bovine Cysticercosis and Human Taeniasis.

### 2.4. Sample Size Determination

Sample size was determined according to Thrusfield (2007) by taking 50% prevalence was considered to calculate based on the following formula

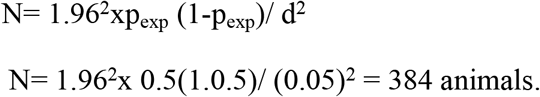

Where, n- Required sample size, Pexp - Expected prevalence, d- Desired absolute precision, usually d is 0.05 at 95% confidence level and 5% expected error.

Hence, the sample size was calculated to be 384. However, sample size was increased to 422 in order to increase the precision of study.

Therefore, the questionnaire survey sample size was calculated by using the formula:

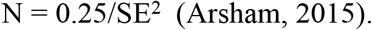

The sample size required for the questionnaire survey as per the above formula was calculated to be 100 but sample size was increased to 110 respondents.

### 2.5. Sampling Technique

Samples in the survey was obtained by systematic random sampling technique using active abattoir slaughter cattle ante-mortem, post mortem examination and questionnaire survey. In selection of slaughter for animal was done by systematic random sampling method where sampling interval (K^th)^ value was obtained by using formulas: - K = N/n.

### 2.6. Study Methodology

For the questionnaire survey, 110 volunteers were selected randomly from the towns. During the active abattoir survey, individual animals were selected using systematic random sampling method sampling interval (K^th)^ value was obtained by using formulas:- K = N/n, during ante mortem examination and post-mortem examination. Before ante-mortem inspection was done, each animal ID is assigned for further follow-up during the post-mortem examination and they were recorded according to their age, sex, and body condition.

#### 2.6.1. Abattoir survey

Active abattoir survey was carried out when routine meat inspection on randomly selected 422 cattle. During pre-slaughtering, identification number, sex, age and origin of animals were recorded. The assessment of body condition, age, sex, breed and their place of origin was determined in ante mortem examination was recorded according to the standard ante-mortem inspection procedures (FAO, 2006). Post mortem examination was made in accordance with the procedures of the Ethiopian guideline by Ministry of Agriculture (MOA, 2015) for the detection of Bovine cysticercosis. The inspection of visceral organs and carcass were done by visualization; palpation and making systematic incisions where necessary for the presence of Cysts and the result was recorded.

#### 2.6.2. Questionnaire Survey

To determine Human Taeniasis and associated risk factors, 110 volunteer respondents were selected using simple random sampling methods based on willingness to participate on questionnaire survey. Selection of respondents for the questionnaire survey was based on random selection of volunteers from selected towns of Silte zone and the interview was conducted face-to-face. The purpose of this interview was to identify the risk factors on their habit of raw meat consumption, frequency of consumption, experience of Taeniasis infection and finding of proglottids in their faces, educational status, age (less than 15-34 years, 34-45 years and greater than 45years years), sex, marital status, knowledge of T. saginata and toilet availability of respondents are registered as possible risk factors (Bebe et al., 2000). The questionnaire was constructed in English and translated to Amharic language.

### 2.7. Data Management and Analysis

Abattoir and questionnaire data collected were entered and coded and preliminary analysis was done using Microsoft Excel work sheet (Microsoft Corporation). Descriptive statistics was carried out to summarize the average prevalence and relative percentage of the disease in each organ. Assessment of association between considered risk factors and prevalence of Cysticercus Bovis was determined by Pearson Chi-square (X^2^). The questionnaire survey collected data was stored by using Epi data software work sheet and analyzed using Statistical Package for Social Science version 22 (SPSS-22) software program. Odds Ratio (OR) was used to determine the effect of different risk factors and logistic regression analysis was used to determine the most significant independent variables and 95% confidence interval (CI) was calculated to assess strength of association of different factors to the occurrence of Taeniasis in Human taeniasis. P- Value less than 0.05 considered significance (p< 0.05).

## 3. Results

### 3.1. Socio-demographic Characteristics

Out of the total of 422 cattle 380 males and 42 females slaughtered cattle inspected ante mortem and postmortem examination data survey in selected Silte zone municipal abattoir during the study period from March 2022 to August 2022. Large numbers of animals were inspected with one meat inspectors due to this Poor meat inspection procedures have been applied in Kilto, Kutery, Kirate and Werabe town municipal abattoir. Also due to small slaughter rooms and weak enforcements of government, backyard slaughtering was practiced. For the ante mortem and postmortem examination, general condition of the animal was observed by visual inspection and age, breed as well as the origin of the animal were recorded.

Out of the total 422 slaughtered animals inspected 152 slaughtered animals 36% positive for C.bovis at postmortem inspection which are 35.7%, 35.92%, 37.96% and 34.11% were Kilto,Kutery,Kirate and Werabe tawon respectively at prevalence recorded.Univariable logistic regression analysis indicated that C. bovis significantly associated with breed, sex and body condition (p < 0.05). In the present study; adult, male, cattle with good body condition and local cattle had significant association with the occurrence of Bovine Cysticercosis than old, female, poor body conditioned and cross breed cattle.There was no statistically significant difference in origin of the animal with the occurrence of Cysticercus Bovis (OR=1.148, P= 0.66) (Table 2).

**Table 2:**
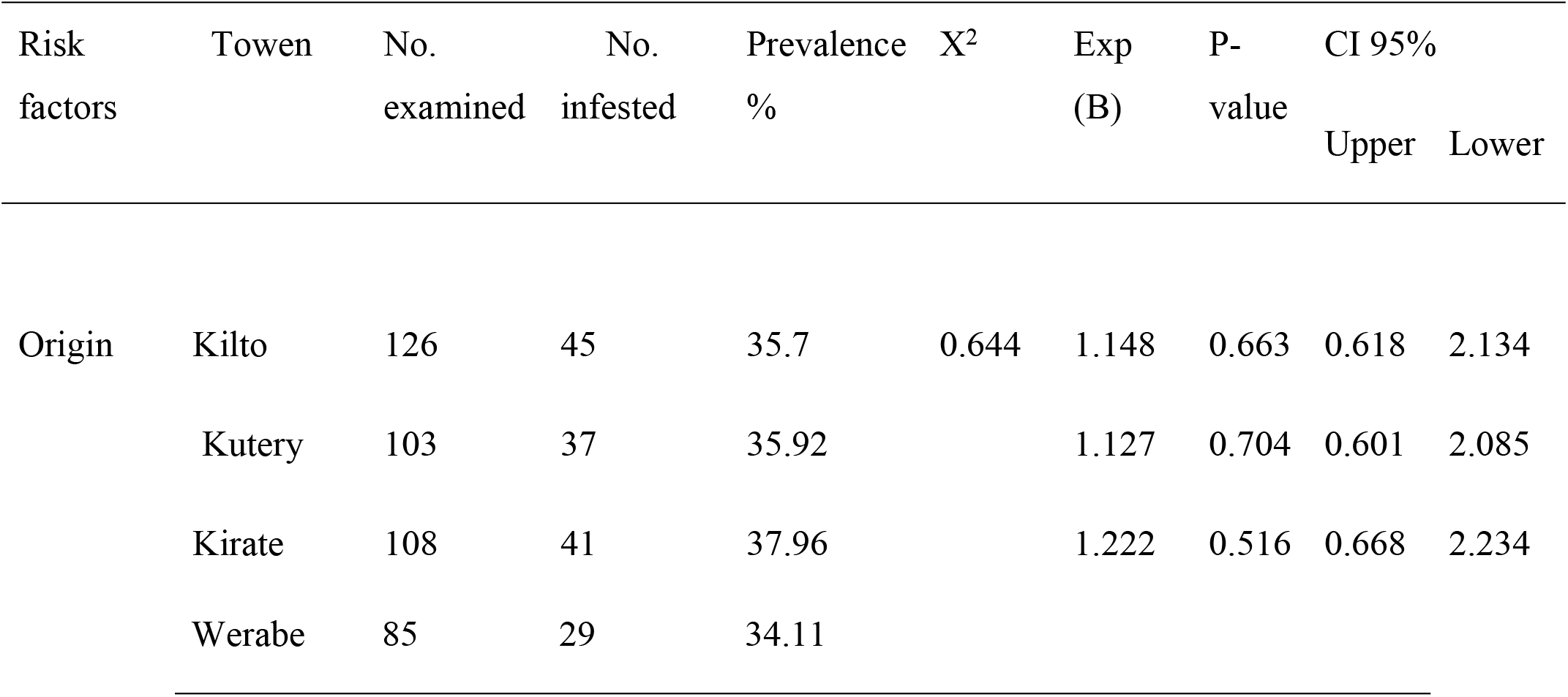
Prevalence of Cysticercus Bovis survey in districts area

For multivariable logistic regression analysis, collinear predictors, namely, sex and breed category were dropped from the model. Sex and breed were found to be collinear with ‘site’. Age non-collinear variables were used to build the final model, and three predictors namely site, sex and body condition were found to be significantly associated with C. bovis (p < 0.05). Adult and good body condition cattle were noted to have higher prevalence than old and poor body condition.Out of the total of 380 males and 42 females inspected female animals 59.52% had higher cysts of C.Bovis than male animals 33.42%.

Although more males than females were examined, the prevalence of infections showed significant difference (OR=3.562, P= 0.000).The total of 273 local and 49 cross breed were inspected and out of these, 145 local 53.11% and 7 cross breed 14.28% positive for C. Bovis with respect to highest prevalence was in local breed than cross breed the prevalence of infections showed significant difference (OR=4.791, P= 0.001). Similarly, statistically significant variation was observed the different categories of body condition (p = 0.002, χ 2 =21.7).

#### 3.1.1. Risk Factors of T. saginata in Human

Among the 110 voluntary interviewed respondents of the residents in the surrounding study data survey in selected Silte zone during the study period from March 2022 to August 2022 at four distinct towen’; namely, Kilto,Kutery,Kirate and Werabe towen participated on different working environments, 63 farmers, 17 students, 25 government employers, and 5 merchants were included interviewed in this study, 71 respondents 64.55% had contracted T. saginata infection. The prevalence on sex basis indicated that 62 males 80.51% were more than 9 females 27.27%; the analysis indicated that males were higher risk than females for Taenia saginata infections. Among the respondents age groups of 34-45 years old 86.66% had relatively higher infection rates than 15-34 years 38%. Univariable logistic regression analysis indicated that T. saginata significantly associated with occupation (civil servant, Farmer, student and merchant), sex, educational status (Illiterate, Primary, Secondary and High-level), age, raw beef consumption, habit of raw beef consumption and form of raw beef (p < 0.05). Age is collinear with occupation, marital status, and educational status and meat sources. In the present study butchers and food related merchants, male, Married, elementary, greater than 18years age raw beef consumer as Kurt, respondents that bought beef other than butchers and those that had no awareness about T. saginata have been positively correlated with the exposure of T. saginata relatively to civil servants and drivers, female, single, other categories of educational status (Primary, Secondary and High-level), less than 18 years age raw beef non consumer, respondents had awareness about T. saginata and bought raw beef from butchers.

Associated risk factors, age groups, gender groups, educational status, occupation,frequency of meat consumption, presence or absence of the latrine, and knowledge about the disease showed statistically significant difference (p<0.05) in the prevalence of Human Taeniasis in this study. Eating raw meat is usually traditional and cultural practice in the study area. Among interviewed, 54 respondents 87.09% had the practice of consuming raw meat and from these 64.55% had contracted the disease. The statistical analysis of the raw meat consumption and Taeniosis interaction was statically highly significant (χ2=10.443 and p=0.003).But no statistical significant association (p>0.05) was observed in the prevalence of Taeniasis between marital status and Meat source (Table 5).

**Table 3:**
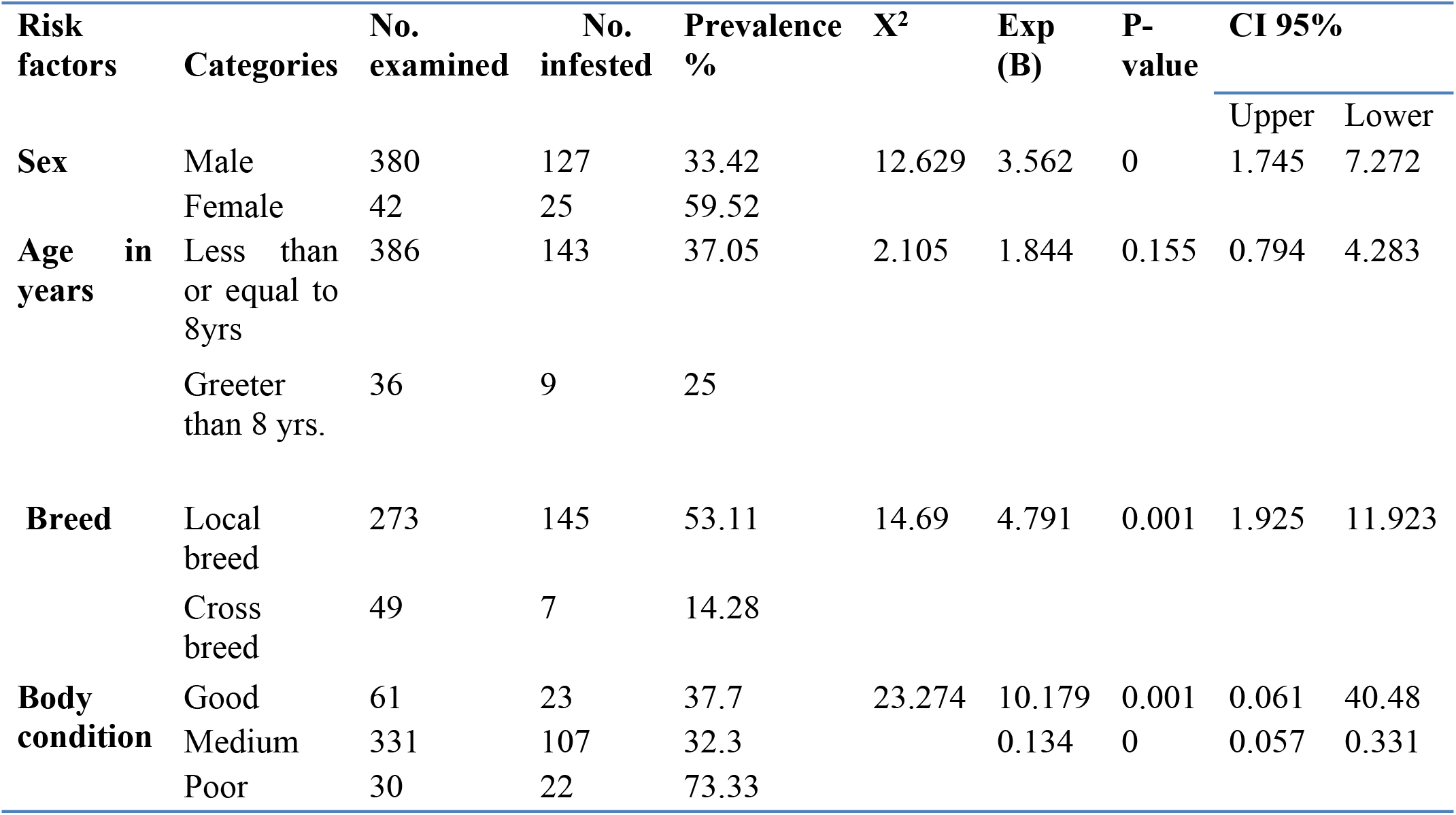
Prevalence of Cysticercus Bovis and associated risk factors

**Table 4:**
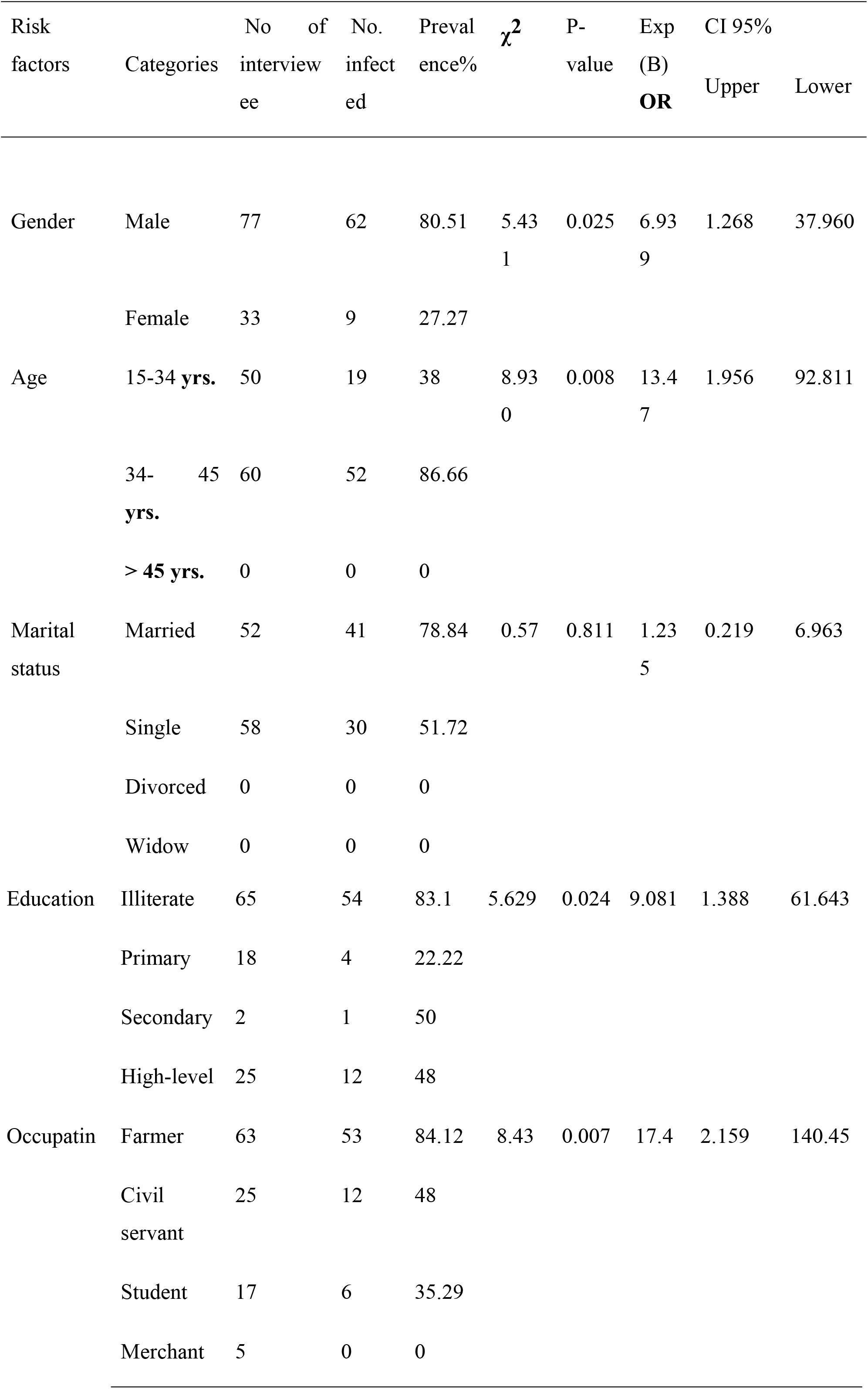
Multivariable logistic regression model for potential predictors of T. saginata

**Table 5:**
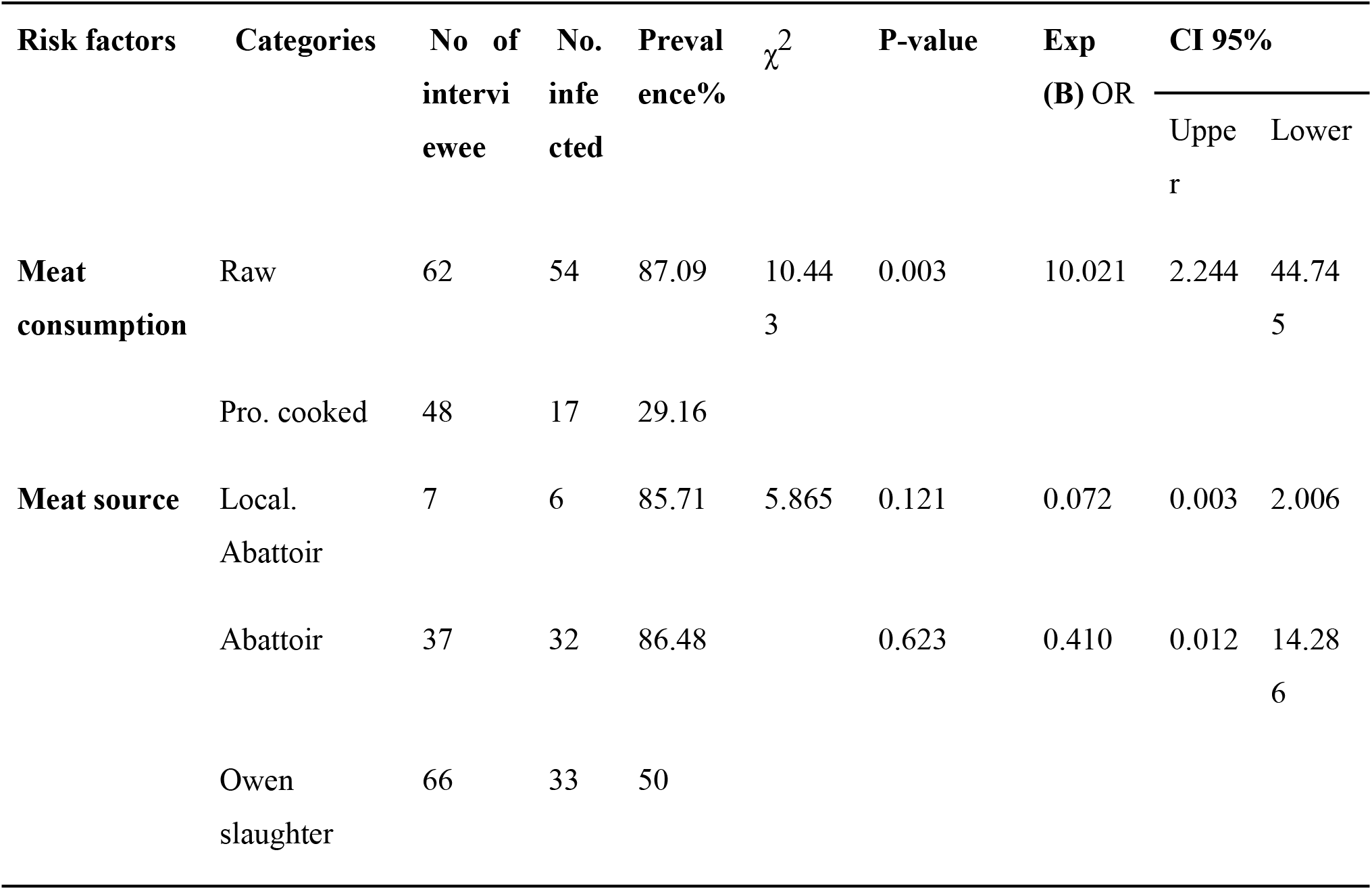
Prevalence of human Taeniasis based on occupational status

#### 3.1.2. Anatomical distribution of cysts

Masseter muscles, tongue, triceps muscles, shoulder muscle, liver and heart organs that were inspected for the presence or absence of Cysticercus Bovis. The anatomical distributions of Cysticercus in result revealed that shoulder muscle was the most frequently infected organ with a prevalence of 28.44%, followed by tongue 20.79%, tricepsbrachi muscle 18.35%, masseter muscle 17.74%, heart 12.84%, and Liver 1.83% (Table4).

## 4. Discussion

### 4.1. Abattoirs survey of Bovine Cysticercosis

Taeniasis/Cysticercosis occurs most commonly in the environments characterized by poor sanitation, primitive livestock husbandry practice, inadequate meat inspection and control. Bovine cysticercosis does not cause much morbidity or mortality among cattle, but it does cause serious economic problems in the endemic areas due to the condemnation of meat or down grading of carcasses (Gizaw and Timetiowos, 2021).

Out of the total 422 slaughtered animals inspected 152 slaughtered animals 36% positive for C. bovis at postmortem inspection. Out of this the prevalence recorded at 35.7%, 35.92%, 37.96% and 34.11% were Kilto,Kutery,Kirate and Werabe towen respectively. The present finding is similar to reports from different parts of Ethiopia, such as,30% from different abattoirs in the country (Fikire et al., 2012) and 30.7% at Eastern Shoa of Oromia (Abdela et al., 2019).

But the current prevalence was greater than from the findings reported by previous authors in different parts of Ethiopia such as 26.25% in Hawassa (Abunna et al., 2008)and 27.6% in Luna export abattoir in East Showa (Hailu, 2005a), 27.6% in Kombolcha (Endris et al., 2011)18.49% in north western Ethiopia (Kebede et al., 2008),13.3% at Walaita Soddo Southern Ethiopia (Birhanu et al., 2014), 11.3% in Walaita Soddo (Southern Ethiopia) (Alemayehu et al., 2009), and 8.6% in Halaba Kulito Towen (Abdul-Aziz et al., 2016) However, the current prevalence was lower than from the findings reported by previous authors in different parts of Ethiopia such as 48.9%, around in Debre Markos abattoir (Regassa et al., 2009) and in Hawassa (Regassa et al., 2010) reported extremely high results (52.69%). This may be due to place to place and also reflects the expertise of meat inspectors in experienced meat inspectors could most likely miss out quite number of viable cysticerci, which blend for with the pinkish-red color of the meat and be passed for human consumption (Adugna et al., 2013).

The present finding on the prevalence of C. bovis is in agreement with earlier reports from African countries, such as 20% in Senegal, 27% in Tanzania (Onyargo et al., 2011; Over et al., 2013), 6.2% in Namibia (Kumba et al., 2010), and (Opara et al., 2012)have reported comparable prevalence of 26.2% from slaughter animals in Nigeria. Conversely, lower prevalence was reported from developed countries, such as 0.26% in Croatia (Zivkovic et al., 2010), 0.48 to 1.08% in Germany (Abuseir et al., 2012) and 0.9% in Cuba (Suarez et al., 2013). But lower than the findings of other authors such as 72.2% prevalence in Nigeria by (Ikpeze et al., 2008) and 38 to 62% in Kenya (Onyargo et al., 2011; Over et al., 2013).

T.saginata/cysticercosis has more public health and economic significance in developing countries like Ethiopia compared with developed countries. Problems associated with poor sanitary infrastructure, low awareness and improper disposal of sewage are major factors for higher prevalence of cysticercosis in developing countries (Gebratsadik et al., 2016). Another possible reason for variation in prevalence may be due to difference in sample size, status of the people in the environment especially related to experience and appropriate use of toilet, habit of the community feeding raw and undercooked meat consumption.

The current study revealed that there was statistically significant between C. bovis infection and breed (local and crossbred) of animals (p = 0.001, χ 2 =14.690). One possible explanation for this significant difference between local and crossbred of cattle in the study area cross breed animals are kept indoor, they are fed only with feed guaranteed free from cysticerci eggs (this means that no feed from pastures or crops can be used, unless treated) and less exposed to human excreta than local breed. This finding is in agreement with (Jemal et al., 2011) reported that the existence of difference in geographical isolates of the parasite and in the breed of cattle as a possible factor affecting the distribution and prevalence of T.saginata cysticercosis.

In this study there was statistically significant difference (P<0.05) between Cysticercosis bovis and sex of the animals. Out of the total of 380 males and 42 females inspected female animals 59.52% had higher cysts of C.bovis than male animals 33.42%. Although more males than females were examined,the prevalence of infections showed significant difference (OR=3.562, P= 0.000). This finding is in agreement with the report of (Meron, 2012) in Jimma and (Worku, 2014) in west Shoa zone Oromia. But disagree with report to no significant association observed between sex (p>0.05), (Kebede et al., 2008), in Addis Ababa, (Gomol et al., 2011) in Jimma and (Meseret et al., 2016) at Kombolcha Elfora.

The current study revealed that there was no statistically significant between C. bovis infection and age of animals (p = 0.156, χ 2 =2.162). This finding is in agreement with reported from (Gomol and Jamal, 2011) at Jimma, (Ibrahim et al., 2012) in Addis Ababa and (Gizaw and Timetiowos, 2021) at Walaita Soddo due to all age group of animals has susceptibility to Taenia saginata eggs. However, present study finding is disagreed with the finding of (Wondimagegnei et al., 2015) showed statistically significant among age of animals due to their different in rank of immunity among age of animals to combat infection.

In the current study high prevalence was statistically significant between C. Bovis infection and body conditioned of animals (p = 0.002, χ 2 =21.683). This finding was higher than the study reported by (Addisu et al., 2015) but lower than the report of (Mesfin et al., 2012b). The reason behind low prevalence in good body condition than medium body condition might be due to the fact that most of the animals slaughtered in the abattoir were brought from fattening systems of the individual farmer, in which animals from such farms were less exposed to eggs of T.saginata.

Regarding the anatomical distribution of the cysts, shoulder muscle showed highest prevalence 28.44%, was followed by tongue 20.79% tricepsbrachi muscle 18.35%, masseter muscle 17.74%, heart 12.84%, and liver 1.8% in order of the proportion of C. bovis affected organ. These preferred predilection sites for the cysts of Cysticercus were similar to earlier reports in Ethiopia (Abshir, 2020), (Meron, 2012),(Hailu, 2005a); (Adugna et al., 2013) and various endemic areas (Anosike, 2001; Cabaret et al., 2002; Opara et al., 2012).

The higher incidence of Cysticercus bovis in some muscles is attributed to increased blood flow due to increased activity, masseter muscle for example was muscle of mastication and similarly shoulder muscle is the most prehensile organ in cattle (Scandrett et al., 2009). The proportion of shoulder muscles affected with C. bovis in this study was 28.44%, which is in agreement with the reports of (Taresa et al., 2011) and (Regassa et al., 2008), 27% and 29.82% respectively and 29.26% at Jimma (Meron, 2012). But the current prevalence was greater than from the findings reported by previous authors in different parts of Ethiopia such 6(1.76%) in Jigjiga Towen (Abshir, 2020).

However, the present finding is lower than the findings of (Megersa et al., 2010) and (Hailu, 2005a), who recoded shoulder cyst proportion of 46.3% and 32% respectively.

#### 4.1.1. Perception and risk factors

The prevalence of human Taeniasis was recorded based on the questionnaire indicated an overall infection rate of 64.55% which was comparable with the reports of (Abunna et al., 2008) who reported 64.2%,(Temesgen,2013) in Shire Indesilassie, Northern Ethiopia (62.5%) and (Tesfaye et al., 2012) in Walaita Soddo (62.5%). But the current prevalence was less than from the findings reported by previous authors in different parts of Ethiopia such (Tariku et al., 2019) 68.6% in Bishoftu, (Hailu, 2005b) in Gondar 79.5% and (Tembo, 2001) in different agro-climatic zones of Ethiopia (89.41%). On the other hand the current prevalence was greater than from the findings reported by previous authors in different parts of Ethiopia such as (Fetene et al., 2014) 58.1% in Jimma town,(Mesfin et al., 2012a) in Addis Ababa (44%) and (Regassa et al., 2009) 50.6% in Walaita Soddo town.

The current study indicates that the ages of respondents have strong association (P=0.008 and χ2 = 8.950) between prevalence of Taenia saginata infection and age distribution. Among the respondents age groups of 34-45 years old 86.66% had relatively higher infection rates than youngest man which finding is in agreement with the previous study of (Meron, 2012) in Jimma, and (Worku, 2014) in west Shoa zone Oromia, have reported that the disease is more common in adults than in children and youth. The possible explanation for the presence of age-wise variation in the prevalence of taeniasis is attributed to frequent raw meat eating habit of adults as compared to those below 15 years (Megersa et al., 2009). The possible suggestion for this could be adult people had the habit of raw meat consumption than the younger population owing to the fact that children are not allowed to consume raw meat and adult individuals can financially afford consuming raw meat “ Kurt or Kitffo” mainly at butcher houses. But disagree (Aragaw et al., 2018) in Kombolcha and (Abdulaziz et al., 2016) at Halaba kulito Towen.

In this study no statistically significant difference were observed between the proportion of taeniasis in marital status although it tends to be higher in single respondents than married respondents (P=0.811 and χ2 = 0.57) which is a similar report with findings of (Meron, 2012a) in Jimma (Worku, 2014) in west Shoa zone Oromia and (Aragaw et al., 2018) in Kombolcha Towen wello. Depending on the marital status, married peoples were more infected than unmarried ones. This might be due to the fact that married peoples have the finance to eat raw beef in the butcher’s house than unmarried peoples.

The current study revealed that raw beef consumers had contracted taeniasis infection more frequently than the non-raw beef consumers (P=0.003 and χ2 = 10.443) which is in lined with the report of (Meron, 2012) in Jmma (Worku, 2014) in west Shoa zone Oromia and (Argaw et al., 2017) in Kombolcha Towen wello. The reason is well known that in the consumption of raw meat the degree of ingesting C.bovis with meat is higher (Geysen et al., 2007). The current study revealed that educational statistically significant difference were observed between the proportion of Taeniasis (P=0.024 and χ2 = 5.629).

The results indicated that uneducated had higher prevalence than those of educated ones. Most of the peoples in the study area were uneducated and with low level of awareness about this helminthic zoonotic disease and low level of education, do not consider Taeniasis as a disease, so that the prevalence of taeniasis was higher in this group than those in higher level of education This report agrees with the finding of (Meron, 2012a) in Jimma, (Abate, 2014) in west Shoa zone Oromia. But disagree (Aragaw et al., 2018) in Kombolcha and (Abdulaziz et al., 2016) at Halaba kulito Towen. Also the current study revealed that the farmers had contracted taeniasis than individuals with other occupational status. This difference might be from low level of awareness in the illiterate and farmers than literate individuals and other occupational status. The other reason for reporting high prevalence of taeniasis in the farmer community is that most of Ethiopian farmers are illiterate and from rural area where environmental hygiene is low and backyard.

In this study the interaction between sex and the prevalence of Taenia saginata was slightly statistically significant (χ 2, 5.431; p=0.025). In agreement with the current results, previous questioner based study in Ethiopia showed that males were at higher risk for T. saginata infection compared with females (Andualem et al., 2017; Lielt et al., 2015;; Meron, 2012b) in Ethiopia. The reason for this significantly higher prevalence in males may be due to economical rezones and cultural practices in Ethiopia that adult men groups often enjoy raw beef (kurt) consumption in butchers and restaurants then than women, where as a great proportion of women in Ethiopia are house wives and commonly prepare their dishes at home consequently females have lower probability of getting viable cysticerci infection. But disagree (Aragaw et al., 2018) in Kombolcha.

Analysis of the results of the present study demonstrate that there was very strong association between raw meat eaters and infection of Taeniasis (χ 2, 5.431; p=0.025). This is a similar report with findings of (Meron, 2012b) in Jimma (Worku, 2014) in west Shoa zone Oromia and (Aragaw et al., 2018) in Kombolcha Towen Wello. But disagree (Bekele et al.,2015) in Jimma and (Abdulaziz et al., 2016) at Halaba kulito Towen.

Computing the logistic regression for the selected risk factors of Taeniasis revealed that age, sex, educational status of the respondents, raw meat consumption, occupation, and latrine available were found to be important risk factors for Taeniasis (Table, 6).Therefore, continues public education should be provided to avoid consumption of raw meat and encourage the use of latrines and improved standards of human hygiene and backyard slaughtering of cattle should be restricted and slaughterhouse which fulfills the necessary facilities and with qualified meat inspector should be constructed.

**Table 6:** Anatomical distribution Post-mortem inspected organs survey in districts area

| No | Infected<br>organs | carcasses infected with *C. bovis*<br>districts area | Total<br>infected<br>*C. bovis* | Prevalence (%) |
| --- | --- | --- | --- | --- |

|  |  | Kilto | Kirate | Kutery | Werabe |  |  |
| --- | --- | --- | --- | --- | --- | --- | --- |
| **1** | Tongue | 22 | 16 | 12 | 18 | 68 | 20.79% |
| **2** | Masseter<br>muscles | 15 | 16 | 16 | 11 | 58 | 17.74% |
| **3** | Shoulder<br>muscles | 38 | 27 | 18 | 10 | 93 | 28.44% |
| **4** | Heart | 15 | 10 | 11 | 6 | 42 | 12.84% |
| **5** | Liver | 3 | 2 | 1 | 0 | 6 | 1.8% |
| **6** | Triceps<br>muscles | 9 | 19 | 21 | 11 | 60 | 18.35% |
|  | Total | 102 | 90 | 79 | 56 | 327 |  |

## 5. Conclusions

The current study assessed the prevalence of C. Bovis and T. saginata in cattle and human populations, respectively. The prevalence of C. bovis in cattle and T. saginata in humans were high. The abattoir survey evidence of the present investigation showed that C.Bovis is prevalent in cattle slaughtered at selected Silte Zone municipal abattoir. The prevalence of C. Bovis was associated with different risk factors such as age, body condition, sex and origin of animals. The most frequently affected organ with the highest number of cysts was the shoulder muscles followed by by tongue, tricepsbrachi muscle, masseter muscle, heart, and liver in order of the proportion of C. bovis affected organ.

The questionnaire survey finding indicated that the infection rate of Taeniasis was higher in the study area. The widespread distribution of Taenia saginata/ Cysticercus Bovis is associated with several factors including; occupation, educational status, marital status, consumption of raw and undercooked meat, bush defecation and poor waste disposal practice, low level of public awareness and presence of backyard (village) slaughtering practices. Slaughter rooms are small, government enforcements are weak and backyard slaughtering was practiced. Besides, large numbers of animals were inspected with one meat inspector. There was also backyard slaughtering practice which could be considered as the contributing factor for zoonosis. Finally, the finding of the present study reflects the zoonotic and economic impact of the disease which needs serious attention by the various stakeholders to safeguard public health.

**Figer 5.**
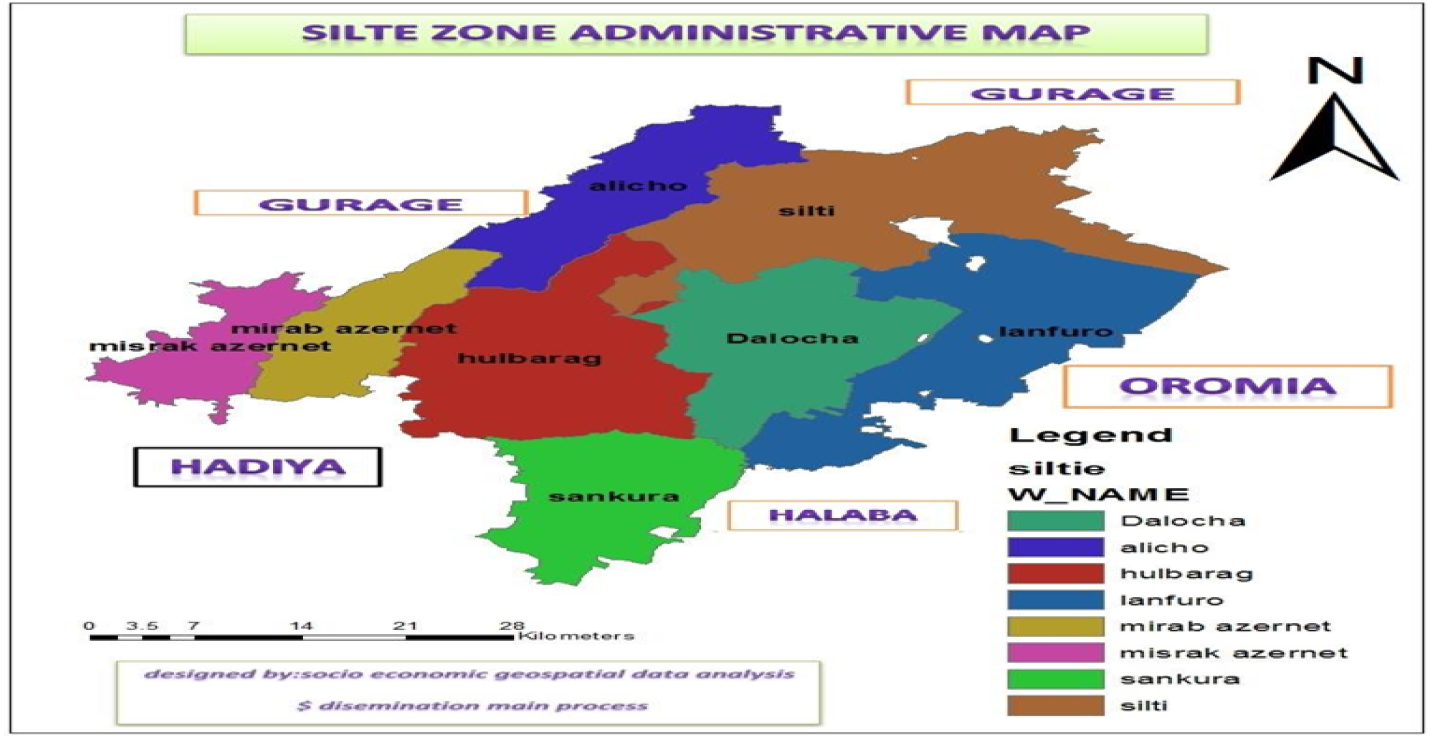
Map of Siltie Zone Administration

## Declaration

### Author contribution statement

### Data availbel statement

Data will be made availbel on request

### Declartion of intersts statmnt

The Authors declare no competing intersts

### Additional information

No additional information is availbel for this paper

